# Event-related and oscillatory signatures of response inhibition: A magnetoencephalography study with subclinical high and low impulsivity adults

**DOI:** 10.1101/2021.03.14.435306

**Authors:** Ainara Jauregi, Hongfang Wang, Stefanie Hassel, Klaus Kessler

## Abstract

Inhibition, the ability to withhold a response or to stop an initiated response, is a necessary cognitive function that can be vulnerable to an impairment. High levels of impulsivity have been shown to impact response inhibition and/or cognitive task performance. The present study investigated the spectral and spatio-temporal dynamics of response inhibition, during a combined go/no-go/stop-signal task, using magnetoencephalography (MEG) in a healthy undergraduate student population. Participants were divided by their level of impulsivity, as assessed by self-report measures, to explore potential differences between high (n=17) and low (n=17) impulsivity groups. Results showed that individuals scoring high on impulsivity failed significantly more NOGO and STOP trials than those scoring low, but no significant differences were found between stop-signal reaction times. During NOGO and STOP conditions, high impulsivity individuals showed significantly smaller M1 components in posterior regions, which could suggest an attentional processing deficit. During NOGO trials, the M2 component was found to be reduced in individuals scoring high, possibly reflecting less pre-motor inhibition efficiency, whereas in STOP trials, the network involved in the stopping process was engaged later in high impulsivity individuals. The high impulsivity group also engaged frontal networks more during the STOP-M3 component only, possibly as a late compensatory process. The lack of response time differences on STOP trials could indicate that compensation was effective to some degree (at the expense of higher error rates). Decreased frontal delta and theta band power was observed in high impulsivity individuals, suggesting a possible deficit in frontal pathways involved in motor suppression, however, unexpectedly, increased delta and theta band power in central and posterior sensors was also observed, which could be indicative of an increased effort to compensate for frontal deficits. Individuals scoring highly also showed decreased alpha power in frontal sensors, suggesting decreased inhibitory processing, along with reduced alpha suppression in posterior regions, reflecting reduced cue processing. These results provide evidence for how personality traits, such as impulsivity, relate to differences in the neural correlates of response inhibition.

## 1. Introduction

Inhibition, the ability to withhold a response or to stop an initiated response, is a necessary cognitive function in many aspects of daily life, as it supports behavioural control (Vara et al., 2014). Because this ability develops gradually from early years to adulthood (Williams et al., 1999), it can be vulnerable to impairments (Rubia et al., 2001), which might negatively influence the daily life of healthy adults (Bari and Robbins, 2013). For example, high levels of impulsivity have been shown to impact response inhibition (Logan et al., 1997; Jauregi et al., 2018).

Impulsivity can be assessed using self-report measures, such as the Barratt Impulsiveness Scale (BIS-11, Patton and Stanford, 1995). Deficits in response inhibition can be behaviourally investigated using a Go/NoGo task (GNGT), which specifically measures “action restraint”, by including trials where a response has to be supressed compared to trials where a response has to be given. Alternatively, the Stop-Signal task (SST) assesses “action cancellation” (Schachar et al., 2007; Eagle et al., 2008; Swick et al., 2011; Bari and Robbins, 2013) by administering a stop-signal shortly after the initiation of a response in relation to a “go” signal. This distinction is supported by growing evidence from neuroimaging studies (e.g., Swick et al., 2011; Sebastian et al., 2013; Dambacher et al., 2014) showing different, in addition to common, neural patterns of activations when both paradigms are examined. Here, previous findings are summarised from studies using electroencephalography (EEG) and magnetoencephalography (MEG). These methods can measure brain activity directly with millisecond temporal resolution (Groß, 2019), which is ideal for measuring rapid brain processes like response inhibition.

### 1.1. Previous findings on ERPs/ERFs

Event-related potentials (ERPs) and event-related magnetic fields (ERFs) offer precise temporal information about the neurological processes underlying response inhibition. We will briefly review relevant ERP components in order of their timing, i.e., from early to late peaking, and separately for GNGT and SST.

During GNGT, a stronger early negative ERP deflection (N1) has been reported to NOGO trials (NOGO-N1) compared to GO trials, between 100ms and 200ms after stimulus onset (De Jong et al., 1990; Filipovic et al., 2000). This negative deflection is suggested to reflect visual detection (Boehler et al., 2009) and attentional processing of stimuli (Vogel & Luck 2000) or infrequent events (Kenemans, 2015). Larger NOGO-N1 amplitudes have been observed in response to certain cues compared to neutral stimuli, which might reflect the amount of attention paid towards stimuli (Gao et al., 2019). Thus, it could be argued that those with enhanced attention towards a cue would show improved task performance, as better attention might facilitate the subsequent inhibition of a motor response in a response inhibition task. Similarly, a MEG study using the SST showed a larger STOP-M1 amplitude for successful relative to unsuccessful trials, suggesting that heightened processing of the STOP cue is also related to success in stopping a pre-potent response (Boehler et al., 2009). However, an EEG study reported larger STOP-N1 amplitude in high compared to low impulsivity individuals (Dimoska & Johnstone, 2007), whereas others have not found different STOP-N1 effects between groups (Lansbergen et al., 2007).

Using the GNGT, Gao et al. (2019) detected a stronger negative ERP deflection (N2) during NOGO trials (NOGO-N2) than during GO trials, occurring between 140ms and 300ms after stimulus onset and which was proposed as an indicator of response inhibition. Chen et al. (2005) reported that impulsive-violent offenders showed reduced NOGO-N2 compared to controls, while others reported larger and/or shorter latencies in individuals with behavioural issues related to impulsivity (Sehlmeyer et al., 2010; Kreusch et al., 2014; Detandt et al., 2017; Gao et al., 2019) or no significant differences at all (Ruchsow et al., 2008). These divergent results could be explained by differences in the measurement of inhibitory deficits and impulsivity scores, requiring a precise assessment of impulsivity. Similarly, a stronger negative deflection in STOP trials (STOP-N2) compared to GO trials in SST has been reported between 200ms and 370ms over frontal areas in ERP studies (e.g., Ramautar et al. 2004; 2006; Schmajuk et al., 2006; Upton, Enticott, Croft, Cooper, & Fitzgerald, 2010), representing increased inhibitory activity (Schmajuk et al., 2006). Although to our knowledge, no significant results have been reported on the STOP-N2 in non-clinical studies using this task, reduced STOP-N2 amplitude and lower reaction time to stop signals have been reported in clinical populations with impulsive characteristics compared to controls (e.g., Pliszka et al. 2000; Dimoska et al. 2003). Altogether, it could be argued that those with worse task performance might show reduced and/or later N2 during NOGO and STOP trials, representing less efficient response inhibition.

Between 300ms to 600ms after stimulus onset, larger positive ERP amplitudes (P3) in NOGO than in GO trials over frontocentral regions have been reported (NOGO-P3; e.g., Huster et al., 2013; Luijten et al., 2014; Wessel & Aron, 2015), which is thought to represent the evaluation of an outcome (Schmajuk et al., 2006; Righi, Mecacci, & Viggiano, 2009; Sehlmeyer et al., 2010). Previous studies have reported reduced NOGO-P3 amplitude in non-clinical impulsive individuals compared to controls (Ruchsow et al., 2008; Benvenuti et al., 2015; Kaiser et al., 2020). Although the literature seems consistent, larger NOGO-P3 amplitudes in non-clinical populations with impulsive characteristics have also been reported (Dong et al., 2010; Detandt et al., 2017). For the SST, a stronger P3 component in STOP trials (STOP-P3) compared to GO trials has been reported to occur between 370ms and 650ms after the presentation of the STOP cue (e.g., Ramautar et al. 2004). The STOP-P3 might be related to the response selection process (Falkenstein, Hohnsbein, & Hoormann, 1994) or to the evaluation of the inhibitory process (Kok et al., 2004). One study (Shen et al., 2014) has reported reduced STOP-P3 amplitudes in a non-clinical population characterised by impulsivity, while others have observed larger amplitudes (Dimoska & Johnstone, 2007; Lansbergen et al., 2007).

Considering the lack of consistency for any of the reviewed ERP components regarding differences between impulsivity groups typically tested separately on either GNGT or SST, it was important to analyse group differences using both paradigms in the same sample.

### 1.2. Previous findings on oscillatory activity

Time-frequency analyses offer additional insights to that of ERPs/ERFs (Doñamayor et al., 2012), as they provide information on the spectral dynamics implicated in response inhibition, shedding light on processes that are not phase-locked to the stimulus onset (Cavanagh and Frank, 2014; Harper et al., 2014). Studies investigating the GNGT have shown an increase in delta (1-4 Hz) power in NOGO trials compared to GO trials (Harper et al., 2014). Delta has been suggested to be involved in response inhibition and may actually serve as an index of motor inhibition (Kaiser et al., 2019). Increased theta band (4-8 Hz) power, specifically in medial-frontal sensors (Kirmizi-Alsan et al., 2006; Harper et al., 2014) has been reported in NOGO trials compared to GO trials (Beste et al., 2011; Huster et al., 2013; Nakata et al., 2013; Isabella et al., 2015; Mückschel et al., 2017). Increased theta band power has been proposed to be essential for response inhibition as an integrative mechanism of early stimulus detection and response selection processes (Cavanagh and Frank, 2014; Mückschel et al., 2017). Although the literature on spectral analysis of the SST is very limited, a recent EEG study using scalp-wide current source density transformation (CSD), found increased mid-frontocentral theta and frontal delta during STOP conditions (Lockhart et al., 2019). It could be argued that in individuals scoring high on impulsivity and showing worse task performance, delta and theta band power might be reduced in NOGO and STOP conditions, reflecting an impairment in motor inhibition, compared to those scoring low and performing better. Decreased delta and/or theta band power during the GNGT have been consistently reported in clinical and non-clinical populations characterised by impulsivity (e.g., Kamarajan et al., 2004, 2006; Colrain et al., 2011; Pandey et al., 2016; Lopez-Caneda et al., 2017). In some studies, this reduction was significantly different only during NOGO trials (e.g., Kamajaran et al., 2006; Krämer et al., 2009) and across frontal regions, reflecting a deficit in frontoparietal networks recruited during inhibitory processing (Kamarajan et al., 2004; Colrain et al., 2011; Lopez-Caneda et al., 2017).

Increased alpha band (8–12 Hz) power has also been associated with response inhibition during NOGO trials (Nakata et al., 2013). Specifically, significantly decreased alpha power in young individuals at risk for alcoholism (Kamarajan et al., 2006) and in abstinent alcohol-dependent adults (Pandey et al., 2016) compared to control subjects, has been reported. This decrease in alpha power has been suggested to reflect an early attentional deficit that might affect the inhibition process (Pandey et al., 2016). To our knowledge, no significant results in alpha band activity have been reported during the SST, but it could be argued that individuals with impulsive characteristics might show a similar decrease in alpha power to those seen in GNGT studies (e.g., Kamarajan et al., 2006; Pandey et al., 2016).

### 1.3. Current study

In a previous behavioural study (Jauregi et al., 2018), we found significantly worse task performance in both, GNGT and SST, in high compared to low impulsivity individuals as measured by self-reports, in a healthy, young population. We also found that combined impulsivity dimensions (rapid-response and reward-delay impulsivity) provide better assessment of impulsivity than each dimension alone and thus, the criteria to be assigned to either group here was based on previous results (Jauregi et al., 2018). In the current study, we use magnetoencephalography (MEG) to further investigate potential group differences in the spectral and spatio-temporal dynamics of response inhibition in this population using the same tasks (GNGT and SST).

As discussed, there is limited and contradictory data on the spectral and spatio-temporal dynamics of response inhibition. It remains inconclusive whether larger or smaller, faster or slower ERP components and/or higher or lower power in delta, theta and/or alpha bands characterise impulsive individuals. In addition, the current dearth of findings that combine GNGT and SST data from the same individuals further hampers a consistent and generalisable understanding of response inhibition and its expression in high impulsivity groups. By using MEG and a combined GNGT-SST task, potential differences in action restraint and action cancellation could be disentangled and applied to high vs low impulsivity groups, providing novel results and potentially clarifying contradictory findings.

All in all, considering common patterns amongst a majority of previous findings, it is hypothesised that in ERFs, high impulsivity individuals are likely to show reduced NOGO-M1, STOP-M1, NOGO-M2, STOP-M2 and NOGO-M3 amplitudes (M denotes magnetic field) compared to low impulsivity individuals, while also showing group differences in STOP-M3 amplitude. Regarding oscillatory activity, decreased power in delta, theta and alpha bands was hypothesised in high impulsivity individuals compared to low impulsivity individuals as the most likely outcome.

## 2. Methods

### 2.1. Participants

Thirty-eight students from Aston University participated in the study, four participants were excluded from MEG analysis, because of low task performance or noisy data. The final 34 participants (28 females, M*age*=18.94, SD*age*=3.9) were ethnically diverse, consisting of White (53%), Asian (41%) and Black (6%) heritage. Experimental procedures were in accordance with the Declaration of Helsinki and approved by the Aston University Ethics Committee, including consent, in writing, prior to any data collection. Participants received either course credit or £10 for each study visit.

The criteria to be assigned to either group was based on a previous study (Jauregi et al., 2018), which showed that combined impulsivity dimensions (rapid-response and reward-delay impulsivity) provide better assessment of impulsivity than each dimension alone. The high impulsivity group (HI; n=17) consisted of participants scoring within the highest 35th percentile of the BIS-11 motor subscale and above 85% on the BAS-Total scale. The low impulsivity group (LI; n=17) included participants scoring below 35% on the motor scale and moderately on the BAS-Total scale, between 40^th^ and 60^th^ percentiles (see Alloy et al., 2012).

### 2.2. Stimuli

A combined Go/No-Go/Stop-Signal Task (GNGSST) was adapted from Boehler et al., 2009, see Figure 1. It was programmed and administered using EPrime 2.0 Professional (Psychology Software Tools; http://www.pstnet.com/), and visualised on a projection screen located 86cm from participants in the MEG room.

**Figure 1.**
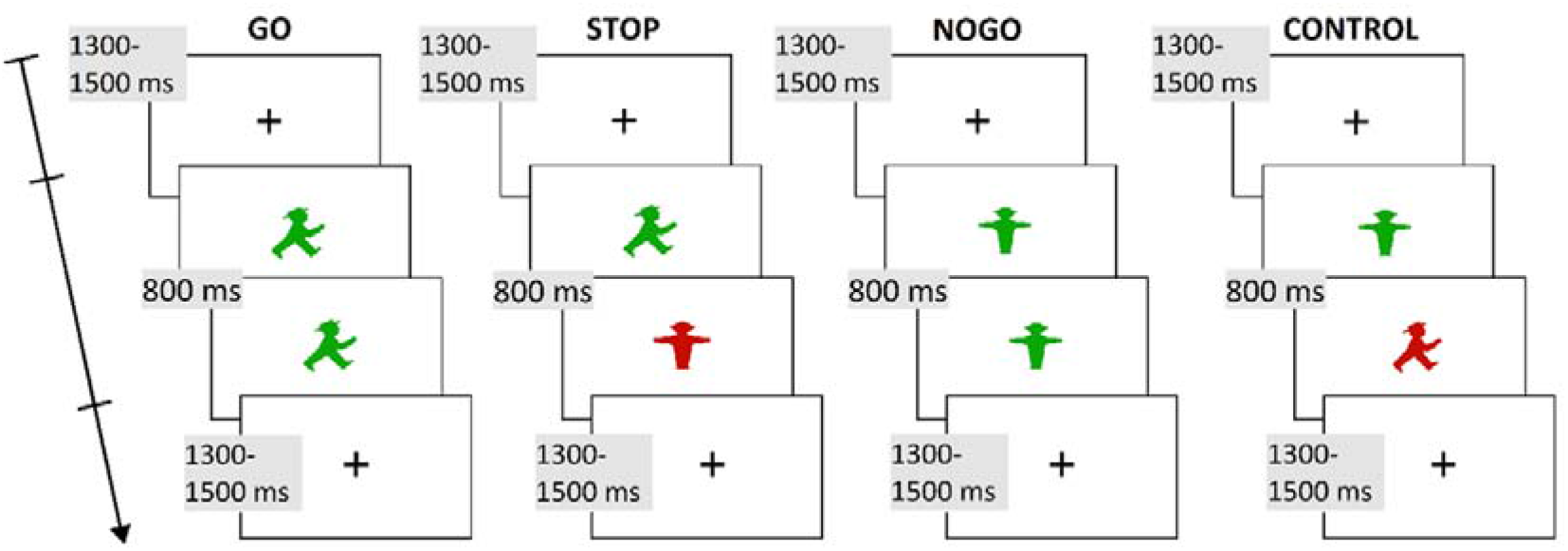
Task description (adapted from Boehler et al., 2009). Trial types in the Go/NoGo-Stop-Signal task. In GO trials the green “GO cue” was presented for 800ms, requiring a button press. In STOP trials the green “GO cue” was substituted by a red “STOP cue” after an SOA set by a staircase-procedure, instructing the participant to cancel the initiated response. Total stimulus duration was 800ms. In NOGO trials a green “NOGO cue” was presented for 800ms, requiring withholding a response. In control-trials the green “NOGO cue” was replaced by a red “GO cue” after an SOA corresponding to those of the STOP trials, which instructed the participant not to respond. Total stimulus duration was 800ms.

Each trial started with a variable fixation period between 1300ms and 1500ms randomised across trials. This was followed by the presentation of one of two possible traffic light symbols: a green “GO-cue”, requiring a button press with the index finger, and a green “NOGO-cue”, requiring withholding a response. Each trial ended when a response was given or when no response was given within 800ms since target presentation. In other trials, the green GO-cue was followed by a red STOP-cue, instructing participants to cancel an initiated response. The stop signal appeared after a stimulus onset asynchrony (SOA) set by a staircase-procedure, in which the SOA was increased 50ms after an unsuccessful stop trial and reduced 50ms after a successful stop trial. The initial SOA was always 200ms and the total duration of the trial always 800ms. In Control-trials, the green NOGO-cue was followed by a red GO-cue, which instructed the participant not to respond, the red GO-cue was presented after an SOA corresponding to those of the STOP trials, while the total duration of the trial was always 800ms. Three blocks were presented, each consisting of 200 trials. The total 600 trials comprised 360 GO-trials (60% of trials), 120 STOP-trials (20%), 60 NOGO-trials (10%) and 60 Control-trials (10%). The variables of interest were the reaction time to the GO-cue, accuracy in GO trials, accuracy in NOGO trials, accuracy in STOP trials and the stop-signal reaction time (SS-RT). The SS-RT, the time that an individual needs to inhibit an already initiated response, can be estimated by subtracting the mean stop-signal delay from the mean GO trial reaction time, a procedure following that of Logan et al. (1997).

### 2.3. Questionnaires

#### Barratt Impulsiveness Scale

The Barratt Impulsiveness Scale (BIS-11; Patton, Stanford, & Barratt, 1995) is a well-validated and reliable measure of trait impulsivity (Stanford et al., 2009). It consists of 30 items, comprising three subscales: non-planning, motor, attentional; rated on a four-point Likert scale (1=rarely/never, 2=occasionally, 3=often, 4=almost always/always). High scores in the sum of all subscales indicate high levels of trait impulsivity as a heterogeneous concept. High scores in each subscale indicate which impulsivity components have a heavier weight.

#### Behavioural Inhibition System/Behavioural Activation System scales

The Behavioural Inhibition System/Behavioural Activation System scales (BIS/BAS; Carver & White, 1994) consist of 20 items, rated on a 4-point Likert-type scale (1=strongly disagree, 4=strongly agree) and comprise three BAS subscales (reward responsiveness, drive and fun seeking) and one BIS subscale (reactions to the expectation of punishment). These scales assess participants’ sensitivity of the behavioural approach system (BAS) and the behavioural inhibition system (BIS) to positive and negative cues. High scores indicate high sensitivity to the BAS or BIS systems.

### 2.4. Data acquisition and analysis

Independent t-tests were conducted to compare the behavioural variables, reaction times (RTs) and accuracy, between the high impulsivity and low impulsivity groups, using SPSS (Version 22.0; SPSS Inc., Chicago, IL, United States). MEG data were acquired using a 306-channel Neuromag MEG scanner (Vectorview, Elekta, Finland) with 204 planar gradiometers and 102 magnetometers, in a magnetically shielded room and at a sampling rate of 1000 Hz. Five head position indicator (HPI) coils were attached to specific sites on the subject’s head for continuous head position tracking and a digital pen, the Polhemus Fastrak, was passed over the scalp to generate a digitised version of the participant’s head. During a subsequent MRI session, a structural T1 scan was obtained for each participant using a Siemens MAGNETOM Trio 3T scanner (Siemens, Munich, Germany).

#### MEG pre-processing

Raw MEG data were first pre-processed separately using system-specific Maxfilter software (version 2.0, Elekta-Neuromag) with temporal signal space separation (tSSS) using a .9 correlation (Taulu and Simola, 2006). Using Fieldtrip, data were then band-pass filtered between 0.5-50Hz, band-stop filtered (49.5-50.5Hz) to remove residual 50Hz line noise and segmented into trials of 1600ms, 800ms baseline and 800ms after onset display, as in previous MEG studies (e.g., Nakata et al., 2005; Boehler et al., 2009). Trials containing artefacts were rejected if the trial-by-channel variance was higher than 8×10-23 (as in Seymour et al., 2017). These steps resulted in an average of 280 successful GO trials, 80 successful STOP trials and 50 successful NOGO trials per participant. Considering the results in Boehler et al., 2009 and that the same task was implemented in the current study, a 50% rate of unsuccessful stopping was expected, yet most participants showed a lower rate, see results section for details. This impeded the analysis of HI vs LI unsuccessful trials and thus, only successful HI versus LI STOP-trials were compared.

#### MEG analysis

All analyses were conducted using Fieldtrip, toolbox version 20161024 (Oostenveld et al., 2011) in Matlab 2014b (Mathworks Inc., Natick, MA). An independent component analysis (ICA) was first conducted to correct for cardiac, ocular and movement artefacts. For the analysis of event-related fields, the averaged ERFs, time-locked to stimulus onset, were computed across trials for each participant, for each condition. Then the 204 planar gradients were combined for each of these averages, resulting in 102-channel combined planar gradiometers, and baselined between −100ms and 0ms from stimulus onset. For the frequency analysis, after pre-processing and ICA, a Hanning taper was applied from 1-30 Hz in steps of 1 Hz, in all trials. Each time window was 200ms, in steps of 50ms. Trials were 800ms pre- and post-cue long, resulting in four cycles per time window. Each participant’s trials were averaged separately for each condition, and planar gradiometers were combined as in ERFs analysis. Cluster-based non-parametric permutation testing, the Monte Carlo method, was utilised to compare the ERFs and TFRs between the HI and LI groups, while also controlling for family-wise error (for details, see Maris & Oostenveld, 2007), with 1000 iterations, two-tailed, and with a cluster-level alpha of 0.05. The p-values were calculated from the maximum cluster-level statistic, if the p-values were smaller than the critical alpha-level, then it was concluded that the data in the NOGO and STOP conditions were significantly different. This approach allowed comparing averaged ERFs during NOGO and STOP conditions, and oscillatory activity in delta (1-4 Hz), theta (4-8 Hz) and alpha (8-12 Hz) bands between HI and LI groups, in each condition.

## 3. Results

### 3.1. Behavioural results

Mean reaction times (RT) for correct GO trials were 532.44ms (±54.0ms) for the high (HI) and 515.11ms (±71.4ms) for the low (LI) impulsivity group, showing no significant difference between groups (*t*(32) = 0.80; p = 0.431). Accuracy for GO trials was 97.7% (±0.6) for the HI group and 98.1% (±0.7) for the LI group, again showing no significant difference between groups (*t*(32) = 1.56; p = 0.130). However, accuracy for NOGO trials was 94.9% (±1.6) in the HI group and 96.1% (±0.8) in the LI group, revealing a significant difference between groups (*t*(32) = 2.59; p = 0.014). In STOP trials, overall accuracy was 72.1% (±8.2) and 78.9% (±7.4) in the HI and LI groups, respectively, revealing a significant difference between groups (*t*(32) = 2.53; p = 0.017). Although mean SS-RT was higher in the HI group (259.56ms; ±58.0ms) than in the LI group (239.58ms; ±52.9ms), this difference was not significant (*t*(32) = 1.05; p = 0.302).

### 3.2. Event-related fields (ERFs)

#### Group differences in ERFs in the NOGO condition

The cluster-based permutation testing performed on ERFs in the NOGO condition revealed significant differences between the HI and LI groups at sensor-level, see Figure 2 (left column). A significant difference between groups was found between 120ms and 190ms after presentation of the NOGO cue (NOGO-M1; p = 0.002, see Methods section for details) in parieto-occipital sensors. Here, the HI group showed significantly smaller amplitude than the LI group. Another significant difference between groups was found during the N2 component, between 210ms and 265ms post-cue (NOGO-M2; p = 0.012), in parieto-occipital sensors. Here, the HI group showed significantly smaller amplitude than the LI group. A significant difference between groups was also found between 270ms and 410ms post-cue (NOGO-M2b; p = 0.002) in left parieto-occipital sensors. Again, the HI group showed significantly smaller amplitudes than the LI group.

**Figure 2.**
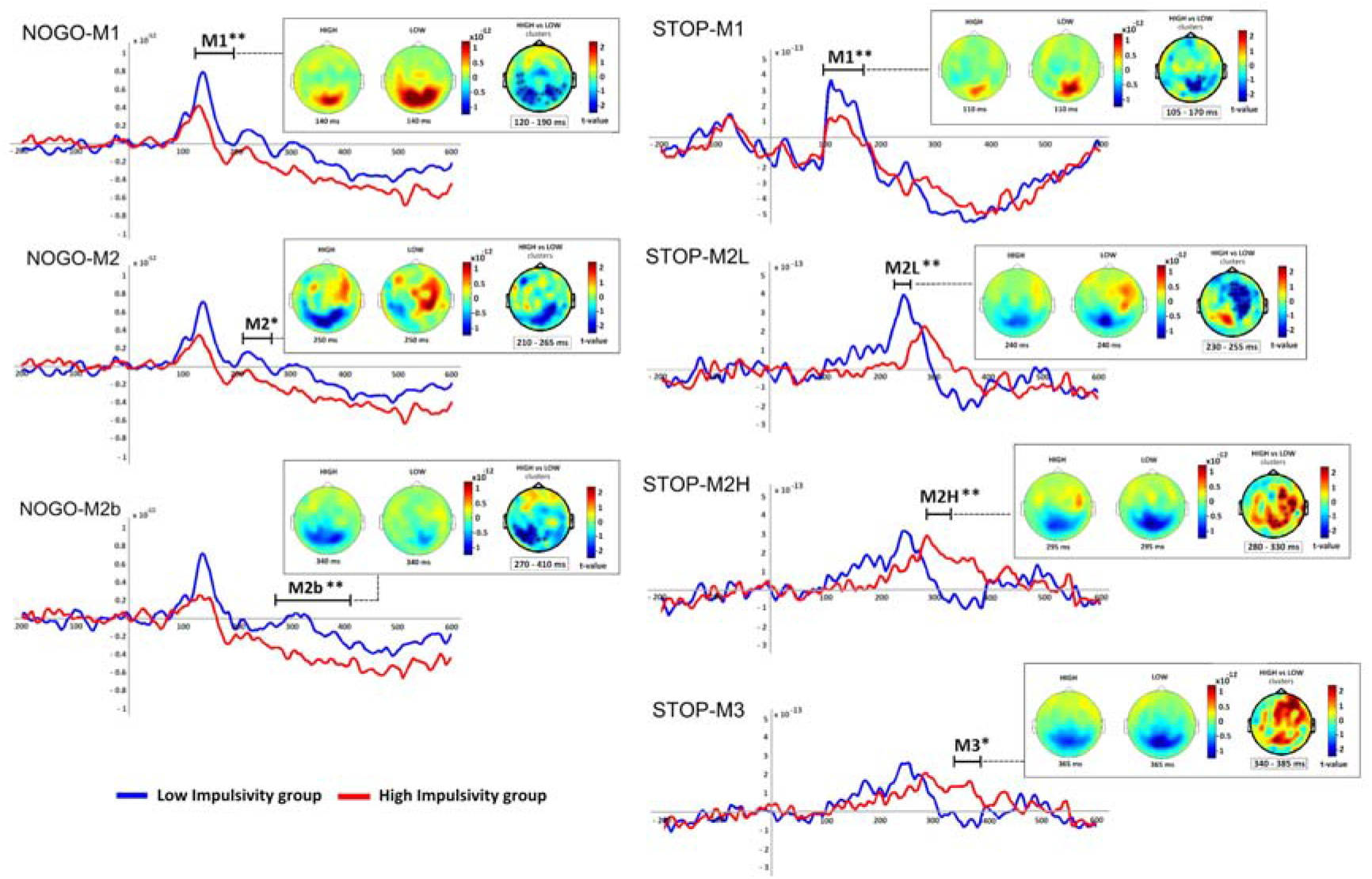
Group differences in event-related fields (ERFs) in NOGO and STOP conditions. Group averages of ERFs during successful NOGO and STOP trials for the HI (red line) and LI (blue line) groups. Group averages were plotted from significant sensors; * p < .05; ** p < .01. Time point 0 denotes onset of the target stimulus presentation. Topography shows each group’s amplitude and the statistical difference between HI and LI groups during the NOGO and STOP conditions. The colour scale represents the *t*-values, hot colours for positive and cool colours for negative values; x indicates significant clusters with p < .05; * indicates significant clusters with p < .01; see Methods section for details.

#### Group differences in ERFs in the STOP condition

The cluster-based permutation testing performed on ERFs in the STOP condition revealed significant differences between the HI and LI groups, see Figure 2 (right column). A significant group difference between 105ms and 170ms after the STOP cue (M1; p = 0.004) was found during the standard time window of the N1 component in parieto-occipital sensors. Here, the HI group showed a significantly smaller amplitude than the LI group. A significant group difference was also found between 230ms and 255ms post-cue. The HI group showed a significant decrease in amplitude compared to the LI group (M2-Low impulsivity group, M2L; p = 0.006) in posterior but also more central anterior sensors. A significant group difference was also found between 280ms and 330ms post-cue. Here, the HI group showed significantly increased amplitude (M2-High impulsivity group, STOP-M2H; p = 0.002) in anterior and right posterior sensors, while amplitude in the LI group was already decreasing (from their STOP-M2 peak), indicating a delayed STOP-M2 in the HI group. The amplitudes of the STOP-M2L and STOP-M2H were compared statistically, but were not significantly different (peak amplitude of NOGO-M2L = 3.6×10e-13; peak amplitude of STOP-M2H = 2.8×10e-13; t(1) = 8.00; p > 0.05). Finally, a significant group difference (p = 0.020) was also found between 340ms and 385ms post-cue in a time window typical for the M3. Here, the HI group showed a significant increase in amplitude compared to the LI group.

#### Theta band activity

Cluster-based permutation tests contrasting HI and LI groups in the NOGO condition with respect to time-averaged power for the frequency range of 4-8Hz, revealed significant differences between groups. As shown in Figure 3 (top half), a significant difference was observed between 0ms and 300ms after stimulus onset (p = 0.009), where the HI group relative to the LI group showed significantly reduced theta power in frontal sensors. A significant difference was also found between 0ms and 652ms after the presentation of the NOGO cue (p = 0.001). Here, HI participants showed significantly higher theta power than LI participants in posterior sensors. In the STOP condition, a significant difference was observed between 0ms and 652ms after stimulus onset (p = 0.001), where the HI group relative to the LI group showed significantly reduced theta power in frontal sensors. A significant difference was also found between 0ms and 652ms after the presentation of the STOP cue (p = 0.001). Here, HI participants showed significantly higher theta power than LI participants in posterior sensors, see Figure 3.

**Figure 3.**
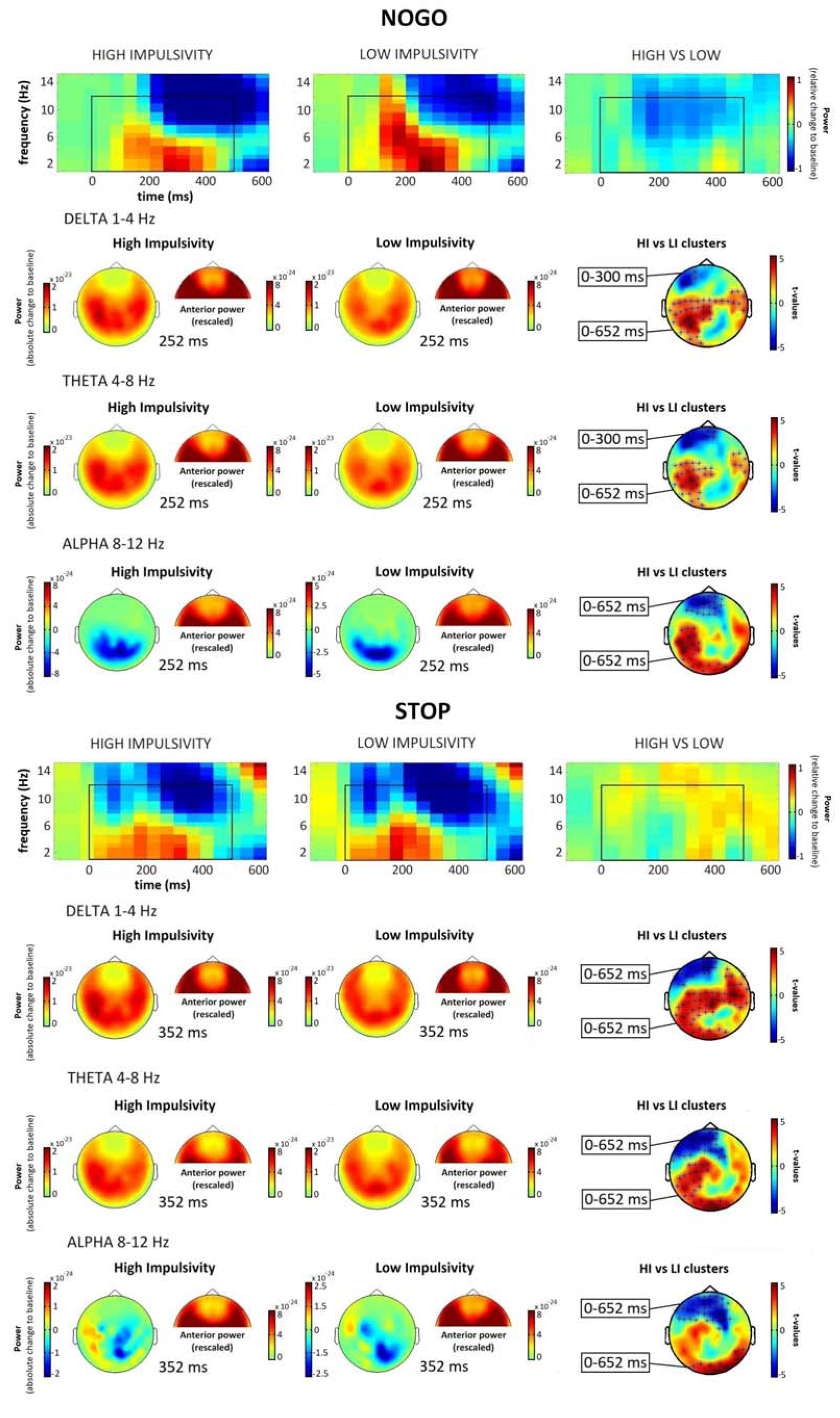
Group differences in time-frequency representations (TFRs) in NOGO and STOP conditions. Baseline corrected TFR power plots for NOGO and STOP trials in HI and LI groups and HI>LI contrast, showing delta, theta, and alpha bands. Y-axis denotes frequency. X-axis denotes time in ms. Colour scale represents logarithmically scaled power change relative to the baseline period, values on the three TFR power plots range from −1 to 1. Below, group average topo-plots in each band power for each condition, colour scale represents baseline-corrected power. On the far right, HI vs LI clusters show statistical differences between HI and LI groups, colour scale represents t-values from cluster-based test permutations, values from −5 to 5. As a result of contrasting the HI against the LI group, hot colours denote positive power difference (HI showing higher power than LI), cool colours denote negative power difference (LI showing higher power than HI). x indicates significant clusters with p < .05. See Methods section for details.

#### Alpha band activity

Cluster-based permutation tests contrasting HI and LI groups in the NOGO condition with respect to time-averaged power for the frequency range of 8-12Hz, revealed significant differences between groups. As shown in Figure 3 (top half), a significant difference was observed between 0ms and 652ms after stimulus onset (p = 0.001), where the HI group relative to the LI group showed significantly reduced alpha power in frontal sensors. A significant difference was also found between 0ms and 652ms after the presentation of the NOGO cue (p = 0.001). Here, HI participants showed significantly higher alpha power than LI participants in posterior sensors. In the STOP condition, a significant difference was observed between 0ms and 652ms after stimulus onset (p = 0.001), where the HI group relative to the LI group showed significantly reduced alpha power in frontal sensors. A significant difference was also found between 0ms and 652ms after the presentation of the STOP cue (p = 0.008). Here, HI participants showed significantly higher alpha power than LI participants in posterior sensors, see Figure 3.

## 4. Discussion

The present study investigated the spectral and spatio-temporal dynamics of response inhibition using MEG, in a healthy undergraduate student population divided by their level of impulsivity as assessed by self-report measures. Impulsive individuals have been reported to show reduced response inhibition, as measured by the number of commission errors in both NOGO and STOP conditions (Jauregi et al., 2018) and longer time to inhibit an already initiated response, the SS-RT, on the Stop-Signal Task (Logan et al., 1997), than controls. Considering our previous results (Jauregi et al., 2018), the HI group was expected to show reduced task performance compared to the LI group, as measured by more commission errors in NOGO and STOP trials. Behavioural results indeed revealed the HI group committed significantly more commission errors in both conditions than the LI group, while the SS-RTs were not significantly different between groups, consistent with our previous findings (Jauregi et al., 2018).

Previous studies investigating differences on response inhibition efficiency measures between high and low impulsivity individuals, as assessed by trait impulsivity questionnaires, have reported contradictory results. Logan et al. (1997) found that the SS-RT was longer in impulsive individuals, as measured by the Eysenck Personality Inventory, which we did not find here. Congruent with our results, Marsh et al. (2002) reported individuals scoring high on impulsivity, as assessed by the Eysenck I7 questionnaire (Eysenck et al., 1985), made significantly more commission errors on the SST than those scoring low. Rodriguez-Fornells et al. (2002), reported no significant differences between high and low Impulsivity individuals, as measured by the Eysenck Personality Inventory (Eysenck & Eysenck, 1964), on the SS-RT, which is consistent with present findings, yet they also reported no significant differences on percentage of correct responses, which differs from current results. Similarly, Lijffijt et al. (2004) did not found significant differences on the SS-RT, when individuals were categorised into high and low impulsivity groups using the Eysenck I7 questionnaire (Eysenck et al., 1985). Dimoska and Johnstone (2007) did not find significant differences between high and low impulsivity individuals, as measured by the Eysenck’s Impulsiveness/Venturesomeness/Empathy (IVE) Questionnaire (1993), on commission errors or SS-RT during the SST. Lijffijt et al. (2004) suggested that these inconsistencies could be explained by sample size. Logan et al. (1997) and Marsh et al. (2002), for example, tested 136 and 86 participants, respectively, whereas Rodriguez-Fornells et al. (2002), Lijffijt et al. (2004) and Dimoska and Johnstone (2007) tested 20, 17 and 40, respectively. They also argued that the method used to assess groups could also explain these differences, as Logan et al. (1997) and Marsh et al. (2002) used a median split analysis (Lijffijt et al., 2004), while the other three studies used the top and lowest scorers on their respective impulsivity questionnaires. In our previous study (Jauregi et al., 2018), which analysed data from 167 participants, we found that only when the two dimensions of trait impulsivity, rapid-response impulsivity and reward-delay impulsivity, were combined to classify participants, significant group differences were observed on the GNGT and SST. Results from the current study, in which 34 individuals were tested, are in line with our previous findings, which makes the sample size hypothesis less plausible, yet strengthening the possibility that the method and questionnaires used when assigning individuals to high or low impulsivity groups might play a crucial role.

In the HI group, compared to the LI group, ERF analysis showed that over posterior sensors M1 amplitude was significantly reduced in NOGO and STOP conditions, that M2 amplitude was reduced in NOGO trials while a later M2 peak was observed in STOP trials, and that, somewhat surprisingly, a larger M3 amplitude was observed in STOP trials. TFR analysis indicated that compared to the LI group the HI group had significantly lower delta, theta, and alpha band power in anterior sensors, and significantly increased power across the three frequency bands in posterior sensors. More specific group differences in ERFs and TFRs during response inhibition and further implications of these findings are discussed below.

### 4.1. Differences in Event-Related Fields (ERFs)

#### NOGO-M1 and STOP-M1

The statistical analysis of event-related fields showed that in the HI group, who showed worse task performance, the amplitude of the NOGO-M1 was significantly smaller in posterior sensors than in the LI group. The NOGO-M1 has been reported to reflect the visual detection (Boehler et al., 2009) and attentional processing of stimuli (Vogel & Luck 2000) or of infrequent events (Kenemans, 2015), suggesting these processes might be reduced in HI individuals. Considering previous and current findings, it could be argued that those with enhanced attention towards the cue show better task performance, as better attention might facilitate the following process, i.e., the inhibition of the motor response. Similarly, the HI group showed significantly reduced STOP-M1 amplitudes in posterior sensors compared to the LI group. Although available data on the SST is very limited in the literature, results from a previous MEG study showed a larger STOP-M1 amplitude for successful than for unsuccessful trials, suggesting that heightened processing of the STOP cue is also related to success in stopping (Boehler et al., 2009). Current findings are consistent with previous results from NOGO conditions, suggesting that reduced processing of NOGO and STOP cues in HI individuals might partly explain their poorer performance in response inhibition tasks.

#### NOGO-M2 and STOP-M2

As expected, a significantly reduced NOGO-M2 amplitude was observed in posterior sensors in individuals scoring high on impulsivity compared to those scoring low. A larger NOGO-N2 amplitude has been reported in participants with better task performance compared to those with reduced performance (Falkenstein et al., 1999; van Boxtel et al., 2001), which is consistent with present findings, and suggests the NOGO-M2/N2 component might serve as a measure of response inhibition efficiency (Schmiedt-Fehr & Basar-Eroglu, 2011). Similarly, a previous non-clinical study showed reduced NOGO-N2 in impulsive-violent offenders compared to controls (Chen et al., 2005). However, others have reported larger amplitudes and/or shorter latencies in individuals with behavioural issues related to impulsivity (Sehlmeyer et al., 2010; Kreusch et al., 2014; Detandt et al., 2017; Gao et al., 2019), or no significant differences at all (Ruchsow et al., 2008). The discrepant results might be explained by differences in the measurement of inhibitory deficits and impulsivity scores. Here, the criteria to be assigned to either group was based on a previous study (Jauregi et al., 2018), which showed that combined impulsivity dimensions (rapid-response and reward-delay impulsivity) provide better assessment of impulsivity than each dimension alone. In the STOP condition, the peak amplitude of the M2 component happened earlier in the LI (STOP-M2L) than in the HI group (STOP-M2H) in central anterior and posterior sensors. Previous studies have shown that the N2 peaks earlier in successful relative to unsuccessful trials (Ramautar et al., 2004; 2006), suggesting that it might be associated with successful inhibition and to represent increased inhibitory activity (Schmajuk et al., 2006). Therefore, it is possible that in individuals scoring high on impulsivity, the network involved in the inhibition process is engaged later. Interestingly, this difference was not observed in NOGO trials, providing further evidence for task-related differences reported in the literature and for the conceptual distinction between “action restraint” and “action cancellation” (e.g., Swick et al., 2011; Sebastian et al., 2013; Dambacher et al., 2014).

#### NOGO-M3 and STOP-M3

High impulsivity individuals were expected to show a smaller amplitude in the NOGO-M3 component, possibly reflecting poor monitoring or evaluation of the inhibitory process (Schmajuk et al., 2006). However, no significant differences between groups were observed in the time period of the M3. The lack of differences is consistent with a previous study (Hoonakker et al., 2017) and suggests that our subclinical HI and LI groups were both monitoring successful motor inhibition to similar degrees. In contrast, in the STOP condition, the amplitude of the M3 component was significantly larger in the HI group in anterior sensors, suggesting this group engaged frontal networks significantly more strongly than the LI group. Individuals scoring high on self-report measures of impulsivity have previously been found to have increased STOP-P3 amplitudes (Dimoska & Johnstone, 2007; Lansbergen et al., 2007). However, others have observed reduced STOP-P3 amplitudes in a non-clinical population characterised by high impulsivity relative to those scoring low (Shen et al., 2014). Current results favour the compensatory strategy proposed in previous studies (Dimoska & Johnstone, 2007; Lansbergen et al., 2007), which suggested that the larger P3 amplitude observed in HI individuals reflects a demand for increased inhibitory effort in this group. The lack of response time differences on STOP trials could further indicate that compensation was effective to some degree, however, at the expense of higher error rates. Present findings therefore dispute the notion that the STOP-P3 reflects the evaluation of the inhibitory process (Kok et al., 2004).

Overall, the results presented here suggest that the impairment found in impulsive individuals might be specifically related to the attentional processing of the stimuli, as reflected by decreased M1, and of the (pre-)motor inhibition process, as indicated by decreased or later M2 in the HI group. The task-related differences found around the M3, suggest the STOP condition might be more sensitive to differences in response inhibition between HI and LI groups than the NOGO condition, further suggesting possible compensatory processing in HI individuals.

### 5.5.2. Differences in oscillatory activity

#### Delta and theta band activity: NOGO and STOP

In the NOGO condition, frequency analysis showed significantly decreased delta and theta power in anterior sensors in the HI compared to the LI group. Decreased anterior delta band power could reflect weaker motor inhibition (Smith et al., 2008), whereas decreased anterior theta possibly reflects a deficit in response selection processes (Cavanagh & Frank, 2014; Mückschel et al., 2017). These suggest that the decreased low frequency (delta and theta band) power observed in the HI group in frontal sensors might reflect a deficit in executive control, specifically, in inhibitory processing. Revealing similar topographies as the NOGO trials, STOP trials also showed delta and theta band power to be significantly decreased in frontal sensors in the HI group compared to the LI group. This is consistent with a recent EEG study showing increased mid-frontocentral theta and increased delta in posterior sensors during STOP trials, in healthy participants (Lockhart et al., 2019). Present findings suggest reduced delta and theta band power in frontal regions might reflect reduced inhibitory processing in high impulsivity individuals during NOGO and STOP trials.

The results of this study are also consistent with those comparing clinical and non-clinical populations during response inhibition tasks. A significant decrease in delta and/or theta power has been reported in individuals characterised by impulsivity, such as young binge drinkers (Lopez-Caneda et al., 2017), compared to controls. It could be argued that the decrease in delta and/or theta power in frontal sensors observed in both NOGO and STOP conditions, possibly reflects a deficit in the frontal part of executive fronto-parietal networks recruited during the suppression of a motor response (Kamarajan et al., 2004; Colrain et al., 2011; Lopez-Caneda et al., 2017). Other studies have proposed that weaker low-frequency oscillatory activity related to inhibitory processing, as observed in delta and theta frequencies, might lead to a predisposition to develop disorders characterised by disinhibition (Kamarajan et al., 2006; Lopez-Caneda et al., 2017). Future studies could conduct a longitudinal follow-up of non-clinical young impulsive adults, to help in understanding how weaker low-frequency activity is related to a possible predisposition to certain disorders.

However, unexpectedly, in both NOGO and STOP conditions, frequency analysis also showed significantly increased delta and theta power in central and posterior sensors in the HI compared to the LI group, which contrasts with previous studies that have consistently found decreased delta and/or theta band power in both clinical and non-clinical populations (e.g., Kamarajan et al., 2004, 2006; Colrain et al., 2011; Pandey et al., 2016; Lopez-Caneda et al., 2017). In some of these studies the reduction was significantly different only across frontal regions in the theta band power (e.g., Kamarajan et al., 2004; Colrain et al., 2011), conforming to our present results, while in others (e.g., Kamarajan et al., 2004, 2006; Colrain et al., 2011; Pandey et al., 2016; Lopez-Caneda et al., 2017), the reduction in delta and/or theta was also found in central and posterior regions, which is contrary to our current findings. This raises the intriguing question what the role of this increased posterior delta/theta signature might be. It is worth noting that the low frequency signatures were very similar across NOGO and SST trials (Fig. 5), possibly suggesting a similar role. Given that an increased M3 was also observed for the HI group in the SST trials and discussed as an indicator of compensatory processing, we propose that the increased delta/theta signature in the HI group could be a further indicator of attempted compensatory processing (Dimoska & Johnstone, 2007; Lansbergen et al., 2007).

#### *Alpha band* activity: *NOGO and STOP*

As in delta and theta activity, significantly reduced alpha band power was observed in anterior sensors during NOGO and STOP trials in the HI compared to the LI group. This finding is consistent with our hypotheses, as an increase in alpha band power in frontal and central areas has been associated with response inhibition during NOGO trials specifically (Nakata et al., 2013). A reduction in alpha activity in frontal regions has also previously been reported in young individuals with impulsive characteristics in the GNGT, such as those at risk for alcoholism (Kamarajan et al., 2006), consistent with current findings in the NOGO condition. To our knowledge, this is the first study that has found a significant reduction of alpha power in anterior sensors during the STOP condition in individuals characterised by high impulsivity.

Results also showed significantly increased alpha power in posterior sensors in HI participants in NOGO and STOP trials, while in LI participants alpha power decreased in the same region. Previous studies, such as Kamarajan et al. (2016) and Pandey et al. (2016), reported reduced slow alpha band power (8-9Hz) in individuals with impulsive characteristics in central and/or posterior regions during NOGO trials, in contrast to current findings. However, studies investigating attentional processing observed that alpha activity increased when suppressing distracting stimuli, whereas alpha activity decreased in regions engaged in the task (Jensen & Mazaheri, 2010, Händel et al., 2011). In an EEG study, adolescents diagnosed with ADHD showed less posterior alpha suppression over occipital electrodes, as reflected by alpha power increase, than typically developed adolescents in a flanker task (Mazaheri et al., 2014). This increase in alpha power, also found in the current study in posterior sensors, has been associated with reduced cue processing (Mazaheri et al., 2014). Here, HI individuals showed increased posterior alpha and decreased anterior alpha power, which could reflect diminished inhibition processing in anterior sensors and a reduced cue processing ability, as reflected by less alpha suppression in posterior sensors.

Altogether, current findings showed decreased low frequency (delta and theta) power in frontal regions, possibly reflecting a deficit in executive control, i.e., inhibitory processing, in high impulsivity individuals, increased delta and theta band power was also observed in central and posterior sensors, which was not expected and could indicate further compensatory processing. Analyses also revealed decreased alpha power in frontal sensors, representing decreased inhibition processing, along with reduced alpha suppression in posterior regions, reflecting reduced cue processing, which appear to result in reduced motor suppression in individuals scoring high on impulsivity compared to those scoring low.

### Conclusions and Limitations

To our knowledge, this is the first MEG study to analyse neural differences between impulsivity groups during NOGO and STOP conditions in the same sample. By doing so, differences in action restraint (NOGO) and action cancellation (STOP) were explored in the same individuals, clarifying previous contradictory findings. Specifically, we observed that in NOGO trials, the HI group showed reduced M2 amplitude, whereas in STOP trials, the peak amplitude of the M2 occurred later in the HI than in the LI group. Surprisingly, differences in the NOGO-M3 component were not observed, yet the amplitude in the STOP-M3 was significantly larger in the HI group. These findings provide further evidence for task-related differences reported previously and might favour the STOP condition as a more effective measure of response inhibition.

One limitation is that not all participants achieved a 50% ratio of successful and unsuccessful number of trials as expected. Considering that the combined task used here had the same conditions and parameters as Boehler et al. (2009), who did find this ratio in a healthy adult sample, this is a surprising result. This suggests a necessity for further improvement of task design. Another methodological limitation is the high number of females tested. Among the thirty-four participants, twenty-eight were females which is not ideal. This was due to Psychology being the most common course studied by participants, in which a higher number of female students are enrolled.

In conclusion, this study showed that impairments in response inhibition in impulsive individuals might be related to reduced attentional processing of NOGO and STOP stimuli. Results suggest that the (pre-)motor inhibition process might be less efficient in impulsive individuals in NOGO trials, whereas in STOP trials, the network involved in the stopping process is engaged later in high impulsivity than in low impulsivity individuals. Findings regarding the M3 suggest the high impulsivity group engaged frontal networks significantly more than the low impulsivity group, possibly as a compensatory strategy, during STOP trials only. In oscillatory activity, current findings suggest that the decrease observed in frontal delta, theta and alpha frequencies in high impulsivity individuals, may indicate that the neural mechanisms underlying cognitive control might be less efficient in these people. High impulsivity individuals also showed significantly reduced posterior alpha suppression compared to low impulsivity individuals, which suggests diminished processing of the NOGO and STOP cues. The results of this study therefore provide evidence for how personality traits, such as impulsivity, relate to differences in the neural correlates of response inhibition.

